# Dependence of Nucleosome Mechanical Stability on DNA Mismatches

**DOI:** 10.1101/2022.11.21.517409

**Authors:** Thuy T. M. Ngo, Bailey Liu, Feng Wang, Aakash Basu, Carl Wu, Taekjip Ha

## Abstract

The organization of nucleosomes into chromatin and their accessibility are shaped by local DNA mechanics. Conversely, nucleosome positions shape genetic variations, which may originate from mismatches during replication and chemical modification of DNA. To investigate how DNA mismatches affect the mechanical stability and the exposure of nucleosomal DNA, we used an optical trap combined with single-molecule FRET and a single-molecule FRET cyclization assay. We found that a single base-pair C-C mismatch enhances DNA bendability and nucleosome mechanical stability for the 601-nucleosome positioning sequence. An increase in force required for DNA unwrapping from the histone core is observed for single base-pair C-C mismatches placed at three tested positions: at the inner turn, at the outer turn, or at the junction of the inner and outer turn of the nucleosome. The results support a model where nucleosomal DNA accessibility is reduced by mismatches, potentially explaining the preferred accumulation of single nucleotide substitutions in the nucleosome core and serving as the source of genetic variation during evolution and cancer progression. Mechanical stability of an intact nucleosome, i.e., mismatch-free, is also dependent on the species as we find that yeast nucleosomes are mechanically less stable and more symmetrical in the outer turn unwrapping compared to Xenopus nucleosomes.

## Introduction

DNA base-base mismatches are generated by nucleotide misincorporation during DNA synthesis, meiotic recombination, somatic recombination between nearly identical repeats, or chemical modification such as hydrolytic deamination of cytosine ^1^. They are also introduced intentionally in some genome editing approaches^2–4^. DNA mismatches, if unrepaired, are sources of genetic variation such as single nucleotide polymorphisms and point mutations which can alter the cellular phenotype and cause dysfunction, diseases, and cancer ^1,5^. DNA mismatches also alter the physical properties of DNA such as local flexibility and conformational heterogeneity ^6–8^.

In eukaryotes, DNA is packaged into a basic unit, the nucleosome, which consists of 147 base pairs (bp) of DNA wrapped around a histone octamer core ^9–11^. In vivo, nucleosomes are regularly arranged along DNA like “beads on a string”, with short linker DNA separating the beads ^9,10^. It has been commonly observed that the rate of genetic variation along the genome is correlated with nucleosome positions.

Although not without an exception^12^, studies have shown that the base substitution rate is higher nearer the center of a nucleosome and increases with increasing nucleosome occupancy^13–20^. One possible explanation for this correlation is that nucleosomes impose a barrier preventing the repair machinery from detecting and repairing a mismatch ^21^ or a bulky DNA adduct induced by ultraviolet^17^, thus leading to substitutions. Currently, it is unknown how substrates for DNA repair such as mismatches and bulky adducts may affect nucleosome mechanical stability and nucleosomal DNA unwrapping, which may affect accessibility of the nucleosomal DNA to the repair machinery.

RNA polymerase II can initiate transcription at 4 pN of hindering force^22^ and its elongation activity continues until it stalls at ∼ 10 pN of hindering force^23,24^. Therefore, the transcription machinery can generate picoNewtons (pN) of force on chromatin as long as both the machinery and the chromatin segment in contact are tethered to stationary objects in the nucleus. Another class of motor protein, chromatin remodeling enzymes, was also shown to induce processive and directional sliding of single nucleosomes when the DNA is under similar amount of tension (∼ 5 pN)^25^. Therefore, measurements of nucleosomes at a few pN of force will expand our knowledge of the physiology roles of nucleosome structure and dynamics.

In an earlier work, we demonstrated a correlation between DNA flexibility and nucleosome stability under tension using the 601 nucleosome positioning sequence ^26^. We showed that the 601 DNA around the histone core can unwrap asymmetrically under tension. One side of the outer DNA turn unwraps at a lower force and the other side unwraps at a higher force. The direction of asymmetry is controlled by the relative DNA flexibility of the two DNA halves flanking the dyad. Unwrapping force is lower for the nucleosomal DNA side with lower flexibility and vice versa. In addition, cytosine modifications that make DNA more flexible made the nucleosome mechanically more stable and vice versa^27^.

Here, we examined the effect of a DNA mismatch on DNA flexibility and nucleosome unwrapping dynamics. We used a single molecule DNA cyclization assay to examine the flexibility of DNA containing a mismatch, and a single-molecule fluorescence-force spectroscopy method to study the effect of mismatch on nucleosome unwrapping dynamics. We also examined the mechanical properties of nucleosomes assembled using yeast and Xenopus histones on intact DNA, i.e. no mismatch, in order to explore the effect of yeast specific histone features^28,29^ on nucleosome mechanical stability.

## Results

### Monitoring nucleosome unwrapping by fluorescence-force spectroscopy

To measure conformational dynamics of the nucleosome in response to external force we used a single-molecule assay that combines fluorescence resonance energy transfer (FRET) with optical tweezers ^30–32^. This assay allows us to use FRET to probe local conformational changes of the nucleosome caused by tension applied by optical tweezers through the two ends of the nucleosomal DNA.

The nucleosome was reconstituted using the nucleosome positioning sequence 601, with or without a C-C mismatch. We designed three DNA constructs 601-R18, 601-R39, and 601-R56 with the mismatches at R18, R39, and R56 positions situated in the middle of the outer turn, at the junction between the outer turn and inner turn, and in the middle of the inner turn, respectively (Fig. 1). Because the distance between the mismatch positions (17, 21 and 38 bp) are not in multiples of 5 bp, they reside in different positions within their own super-helical turn. And the mismatches are at positions where the major groove face toward (R56) or away from (R18, R39) the histone core (R56) so they do not all share the same specific contacts with the histone octamer. Two fluorophores – Cy3 (FRET donor) and Cy5 (FRET acceptor) - were placed in appropriate positions to report on the unwrapping of various sections of nucleosomal DNA through reduction in FRET (Figure 1 and Supplemental Figure S1). The two strands of the DNA construct were separately created by ligation of the component strands (Supplementary Fig. S1) to ensure that the resulting DNA does not contain a nick. The double-stranded construct was then formed by slowly annealing the two purified ligated strands over 3-4 hours. All four DNA constructs (601, 601-R18, 601-R39, and 601-R56) yielded nucleosomes with the same electrophoretic mobility and single molecule FRET value, indicating that the nucleosomes are homogeneously positioned for all four constructs (Supplementary Fig. S1). This is consistent with a previous single nucleotide resolution mapping of dyad position from of a library of mismatches in all possible positions along the 601 sequence or a budding yeast native sequence which showed that a single mismatch (A-A or T-T) does not affect the nucleosome position^33^.

**Figure 1:**
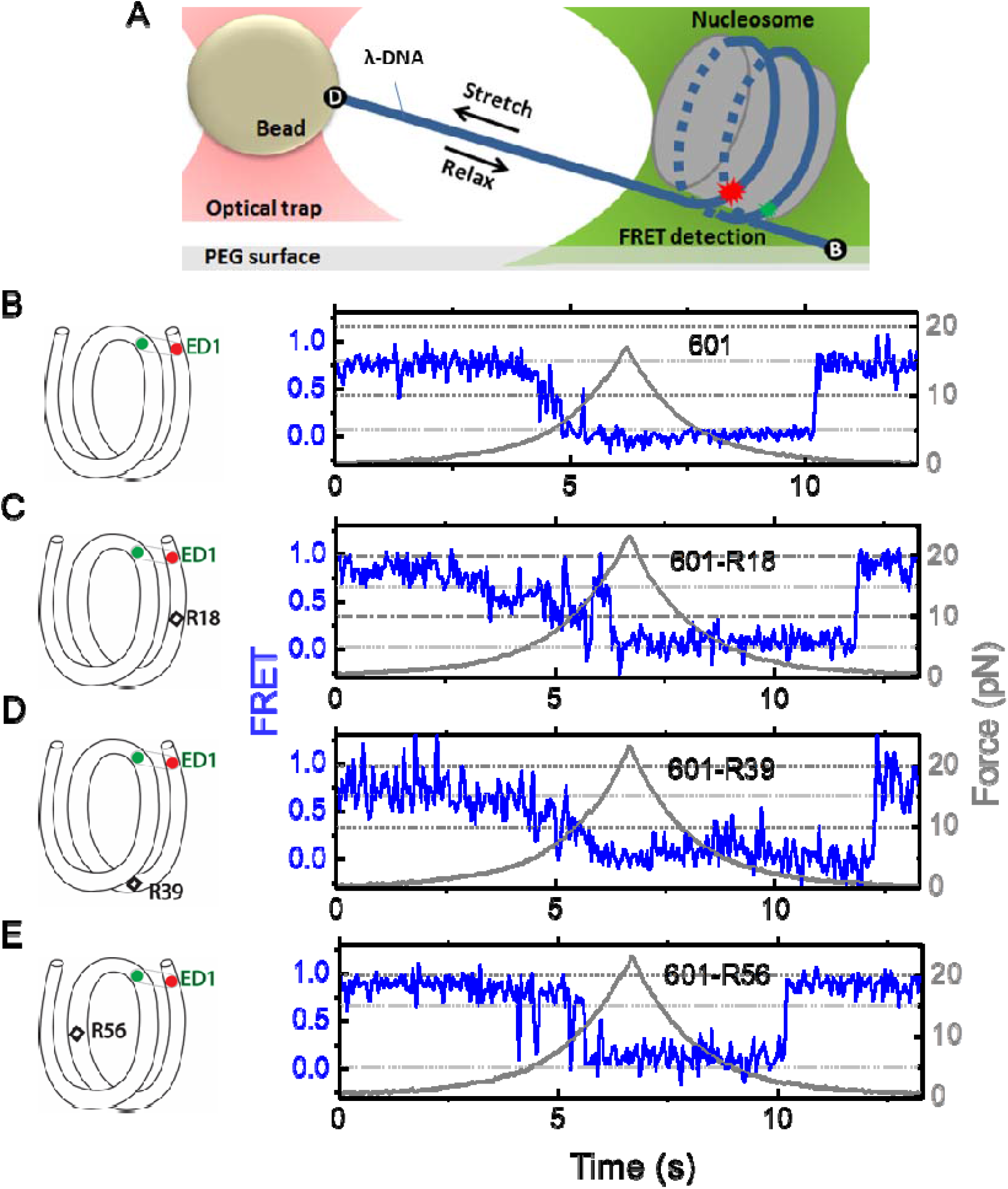
Nucleosome unwrapping measurement. (**A**) Experimental scheme. The red and green stars represent labelled Cy5 (acceptor) and Cy3 (donor) fluorophores, respectively. Biotin, B, and digoxigenin, D, are used to tether the nucleosome-lambda DNA construct to the surface and the bead, respectively. (**B, C, D, E**): Representative stretching traces of the outer turn (ED1) for nucleosomes reconstituted from the 601 sequence (**B**) and from the 601 sequence with containing a mismatch at different positions: on the outer turn (**C**), at the junction of the outer turn and inner turn (**D**) and at the inner turn (**E**). The red and green dots on the DNA bends represent labelled Cy5 and Cy3 fluorophores. The elongated circles enclosing red and green dots represent the ED labeling position. The black diamonds on the DNA bends represent the mismatch position with R18 and R39 on histone=facing minor grooves and R56 on a histone-facing major groove.

In the fluorescence-force spectroscopy assay, a nucleosome was anchored to a polymer-passivated glass surface via biotin-neutravidin linkage on one end of the nucleosomal DNA. The other end of the nucleosomal DNA was attached to a bead held in an optical trap via a λ-DNA tether (Fig. 1A). As previously described ^26^, we attached a pair of donor and acceptor fluorophores to the DNA to probe the unwrapping of nucleosomal DNA. To probe the unwrapping of the outer DNA turn, we constructed DNA with a labeling scheme called ED1 (end-dyad 1) in which the donor is incorporated on the 68^th^ nucleotide from the 5’ end of the top strand (I68) and the acceptor is attached to the 7^th^ nucleotide from the 5’ end of the bottom strand (J7) (Fig. 1A). Upon nucleosome formation, the ED1 probe displayed high FRET due to proximity between the donor and the acceptor. We applied tension to the nucleosomal DNA by moving the piezo stage to which the glass surface attached at a constant speed of 455 nm/s while a focused laser (532 nm) follows the molecule to monitor fluorescence signals. The force increases nonlinearly and the loading rate, i.e. the rate at which the force increases, was approximately in the range of 0.2 pN/s to 6 pN/s, similar to the cellular loading rates for a mechanosensitive membrane receptor ^34^. The force was increased from a low value (typically between 0.4 – 1.0 pN) to a predetermined higher value and then returned to the low value by moving the stage in the opposite direction at the same speed (Fig. 1). We observed a gradual decrease in FRET - corresponding to an increase in the Cy3-Cy5 distance - as the force increases. Upon further increase in force, we observed rapid fluctuations in FRET, followed by a sharp decrease in FRET (Fig. 1), consistent with our previous studies ^26,27^ and a more recent study ^35^ utilizing high resolution optical tweezers with simultaneous smFRET detection. Upon relaxation through gradual decrease in force, the nucleosome reformed as reported via recovery of high FRET but at a lower force than the force at which unwrapping occurred, demonstrating mechanical hysteresis.

### History-dependent mechanical stability of mismatch-containing nucleosomes

Electrophoretic mobility shift analysis and zero-force FRET values did not show a noticeable difference between unmodified nucleosomes and mismatch-containing nucleosomes (Supplementary Fig. S1), consistent with a previous study that showed that, at single nucleotide resolution, a single mismatch does not change the dyad position^33^. However, under perturbation by force, although unmodified nucleosomes showed the same behavior between stretching cycles ^26^, mismatch-containing nucleosomes showed different behaviors between stretching cycles (Fig. 2). The ED1 side of the mismatch containing nucleosomes unwrapped at lower forces for the first few cycles and then at higher forces for subsequent cycles. After relaxation, we observed a general trend of an increase in unwrapping force in the subsequent stretching cycles. One possible explanation for this observation is the re-positioning of the nucleosome such that the mismatch moves toward the dyad, bringing the ED1 probe toward the interior of the nucleosome, as predicted by a previous theoretical model ^36^. According to this model, the nucleosome position is weakly affected by the presence of a flexible lesion on the DNA, but under perturbation by other cellular components which either stiffen the DNA overall or weaken histone binding, the lesion can be made to have a strong preference for the dyad position. In our experiments, applied tension during stretching may act as perturbation which weakens nucleosome binding. When the probes move closer to the dyad in the subsequent stretching cycles, more base pairs of DNA would need to be unwrapped for FRET to decrease, potentially explaining the observed increase in unwrapping force. However, since the FRET values in our DNA construct are not sensitive to the nucleosome position, further experiments with fluorophores conjugated to strategic positions that allow discrimination between different dyad positions^37^ will be required to test this hypothesis.

**Figure 2:**
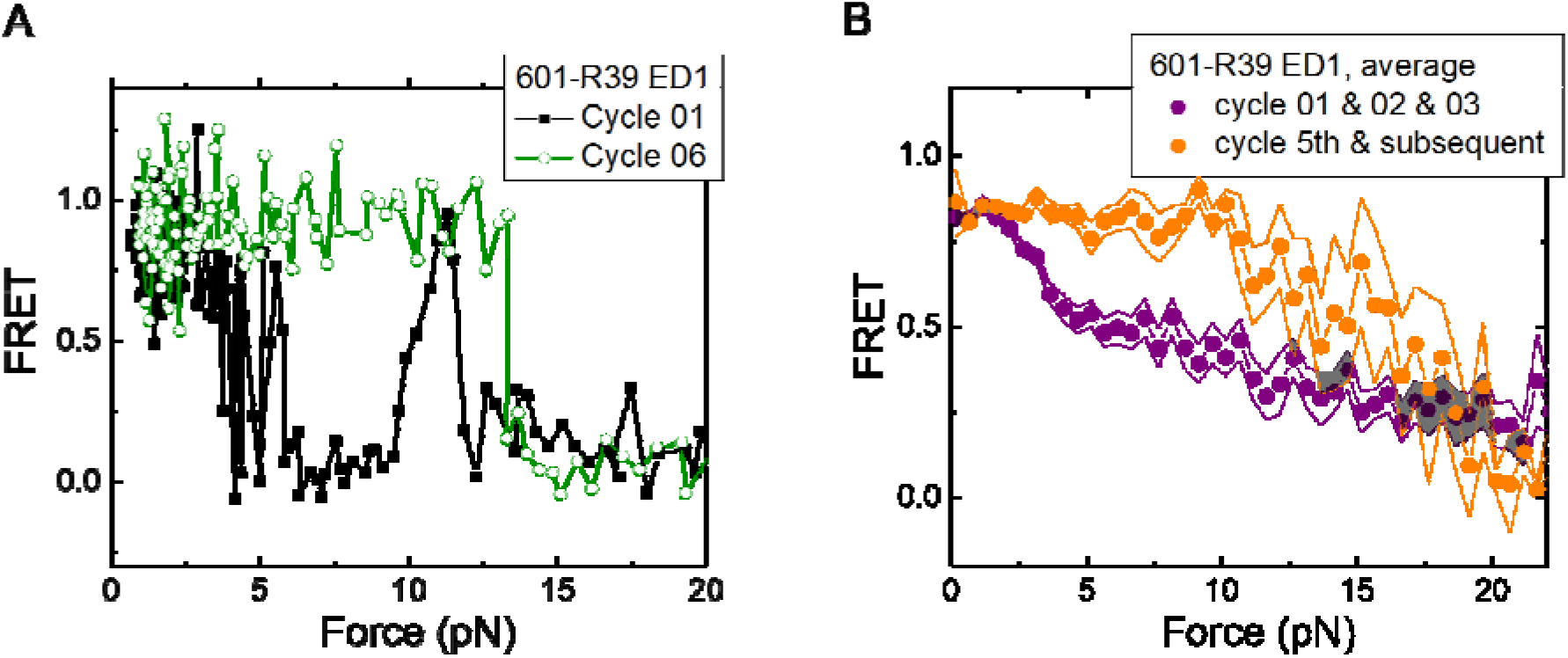
Unwrapping force of mismatch-containing nucleosomes is higher for subsequent stretching cycles. **(A**) Representative single-molecule stretching traces at two stretching cycles from the sample molecule, probe by the ED1 FRET pair in the 601-R18 nucleosome. (**B**) Averaging FRET vs. Force for many molecules at the first three stretching cycles (purple) and the subsequent stretching cycles (orange). Histone proteins were expressed in xenopus. The error bars represent S.D. of n = 25 and 11 traces for the first 3 stretching cycles (purple) and for the cycle 5^th^ and the subsequent stretching cycles (orange), respectively.

### DNA C-C mismatch enhances nucleosome mechanical stability

We compared the FRET versus force curves for constructs containing one C-C mismatch each at three different locations: R18, R39 and R56. We observed similar stretching patterns for the mismatch-containing nucleosomes (601-R18, 601-R39, 601-R56) to that of the 601 nucleosomes. However, the force range where FRET reduced gradually accompanied by fluctuations was wider and extended to higher force for mismatch-containing nucleosomes (Fig. 1 C-E). Because we observed increases in unwrapping forces for the second stretching cycle and beyond for mismatch-containing nucleosomes, we only used the first stretching cycle for comparing unwrapping forces between constructs. The averaged FRET vs. force pattern for 601-R39 showed an increase in unwrapping force for the mismatch-containing nucleosomes (Fig. 3A). The increase in unwrapping force for all three mismatch containing constructs indicates that local flexibility of either the inner turn or the outer turn regulates nucleosome unwrapping (Fig. 4). Next, we probed unwrapping of the nucleosome on the side that does not contain the mismatch for the construct containing a mismatch at the R39 position. In this configuration named ED2 (end-dyad 2), the donor was placed on the inner DNA turn close to the dyad (J58) which is similar to the ED1 construct, and the acceptor was incorporated to the opposite ends (I9) (Fig. 3B). Stretching curves of ED2 nucleosomes formed on the 601 sequence yielded higher unwrapping force compared to the ED1 side as reported previously ^26^. The mismatch construct yielded nearly the same unwrapping pattern as the 601 nucleosome, suggesting the change in local flexibility induced by the mismatch has a strengthening effect against unwrapping only for the side containing the mismatch (the ED1 side).

**Figure 3:**
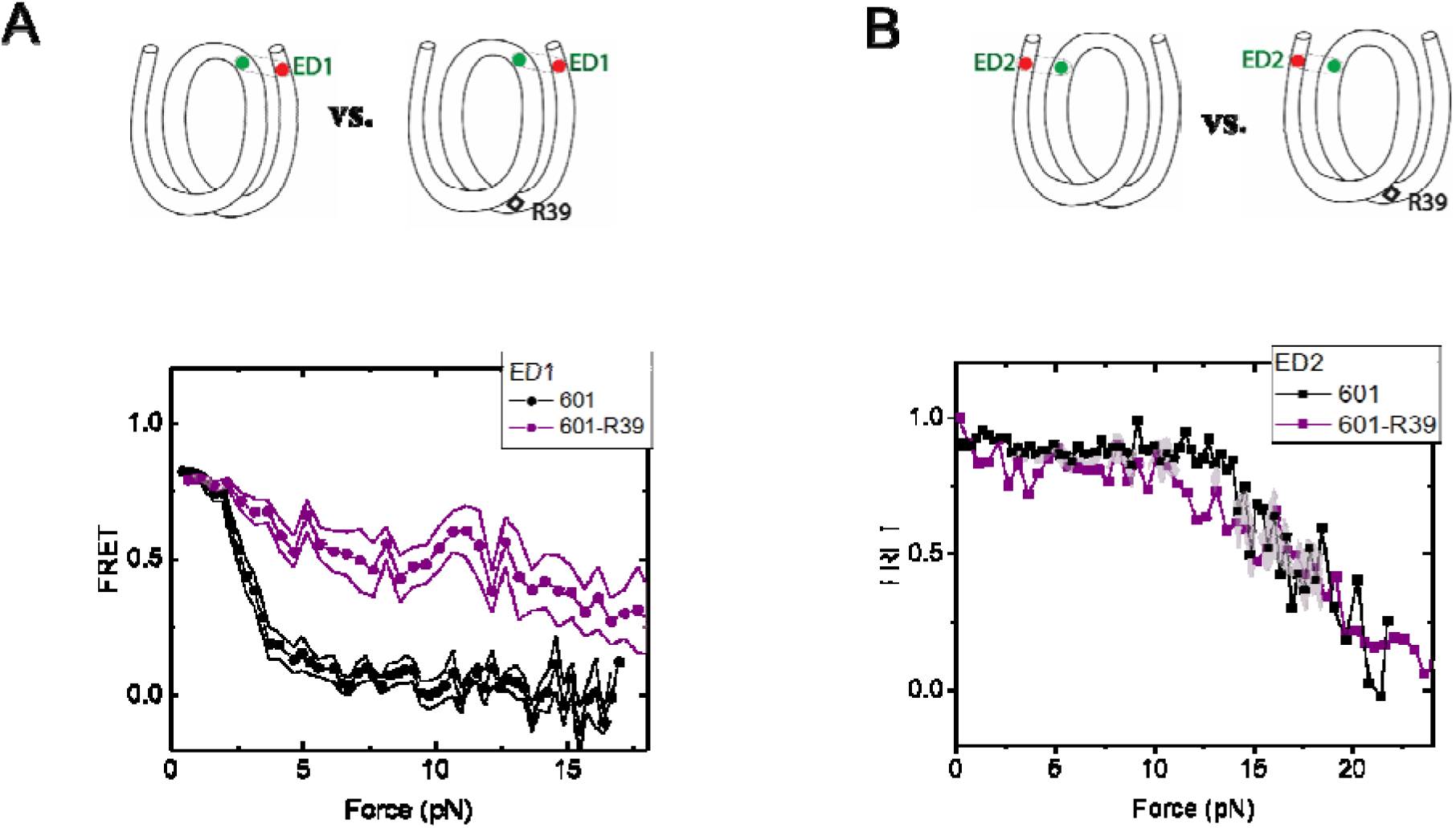
Enhancement of nucleosome mechanical stability by DNA mismatch. Average of FRET vs. Force for ED1 probe (**A**) and ED2 probe (**B**) for the 601 nucleosome (black) and for the first stretching cycle of the mismatch containing nucleosome 601-R39 (purple). Histone proteins were expressed in xenopus. The error bars represent S.D. of n = 25 and 7 for the ED1 probe of the 601 and 601-R39 nucleosomes (**A**) and n = 20 and 39 for the ED2 probe of the 601 and 601-R39 nucleosomes (**B**), respectively.

**Figure 4:**
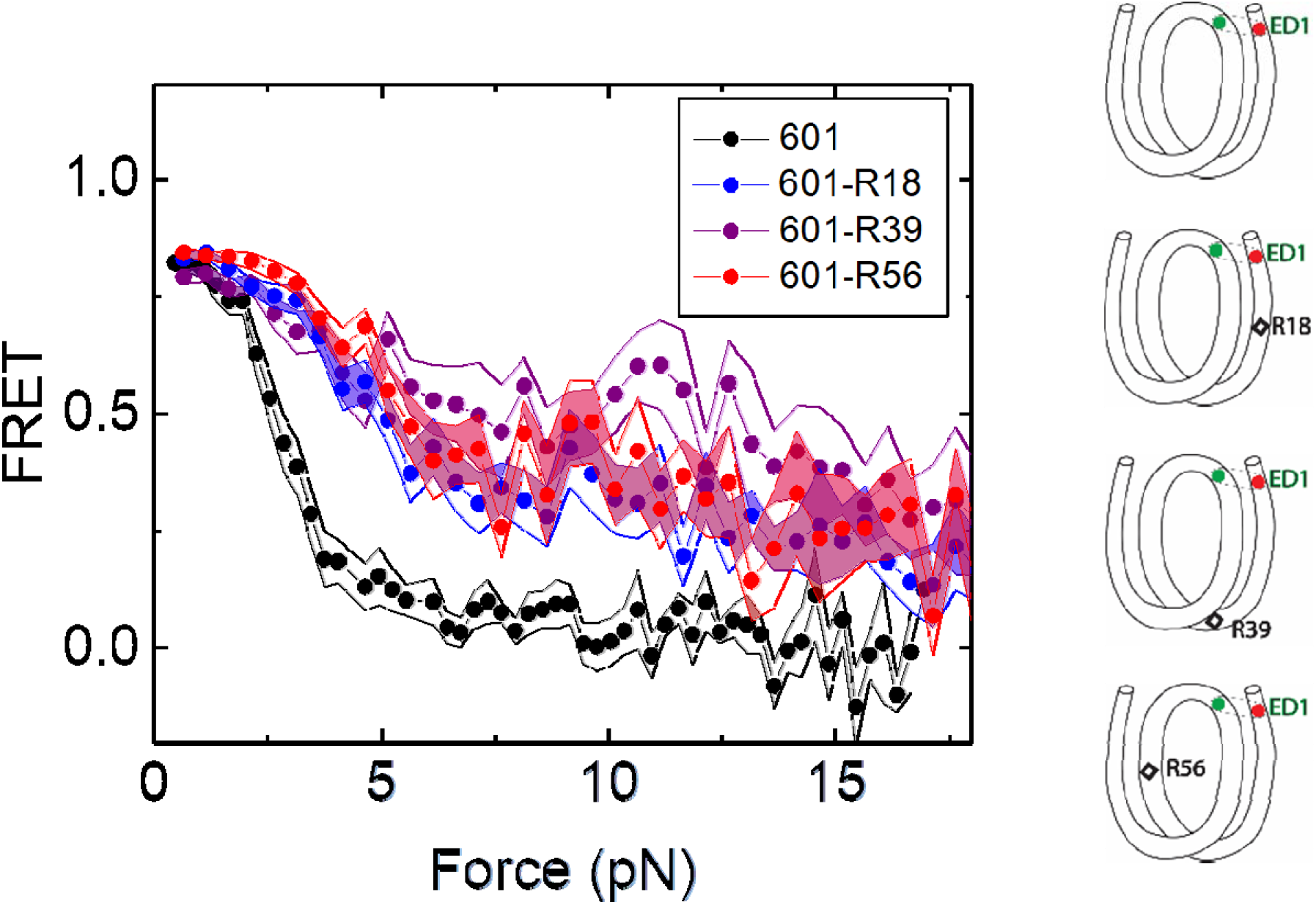
Mismatch position-dependence of nucleosome unwrapping. Average of FRET vs. Force for ED1 probe for the 601 nucleosome (black) and the mismatch-containing nucleosome 601-R39 (purple), 601-R18 (blue) and 601-R56 (red). Histone proteins were expressed in xenopus. The error bars represent S.D. of n = 25, 11, 7 and 10 for the 601, 601-R18, 601-R39 and 601-R56 nucleosomes, respectively.

### DNA C-C mismatch enhances DNA bendability

A single DNA mismatch can cause DNA to deviate from the B-form conformation ^7,8^ and increase DNA flexibility ^6^. A previous study using a DNA buckling assay suggested that C-C is one of the most flexible mismatches ^6^. Therefore, we chose C-C as a representative mismatch to investigate its effect on nucleosome stability. We hypothesized that the stabilization of the nucleosome forming on mismatch containing DNA sequences is caused by its increase in DNA bendability. Therefore, we used a single molecule DNA cyclization assay ^38^ to probe the change in apparent bendability of the right half (RH) of the 601 sequence upon introducing the C-C mismatch. In this assay (Fig. 5), DNA fragments with two 10 nt long 5’ overhangs were immobilized on a microscope slide. A FRET pair (Cy3 and Cy5) was incorporated at the 5’ ends of the overhangs that are complementary to each other, allowing us to detect high FRET when the two overhangs anneal with each other forming a circle. We used smFRET to quantify the fraction of looped molecules versus time after the high salt buffer is introduced in the chamber. The rate of loop formation, which is the inverse of looping time determined from an exponential fitting of loop fraction vs time, was used as a measure of apparent DNA flexibility influenced by a mismatch ^39,40^. The faster the looping occurs, the more flexible the DNA is.

**Figure 5:**
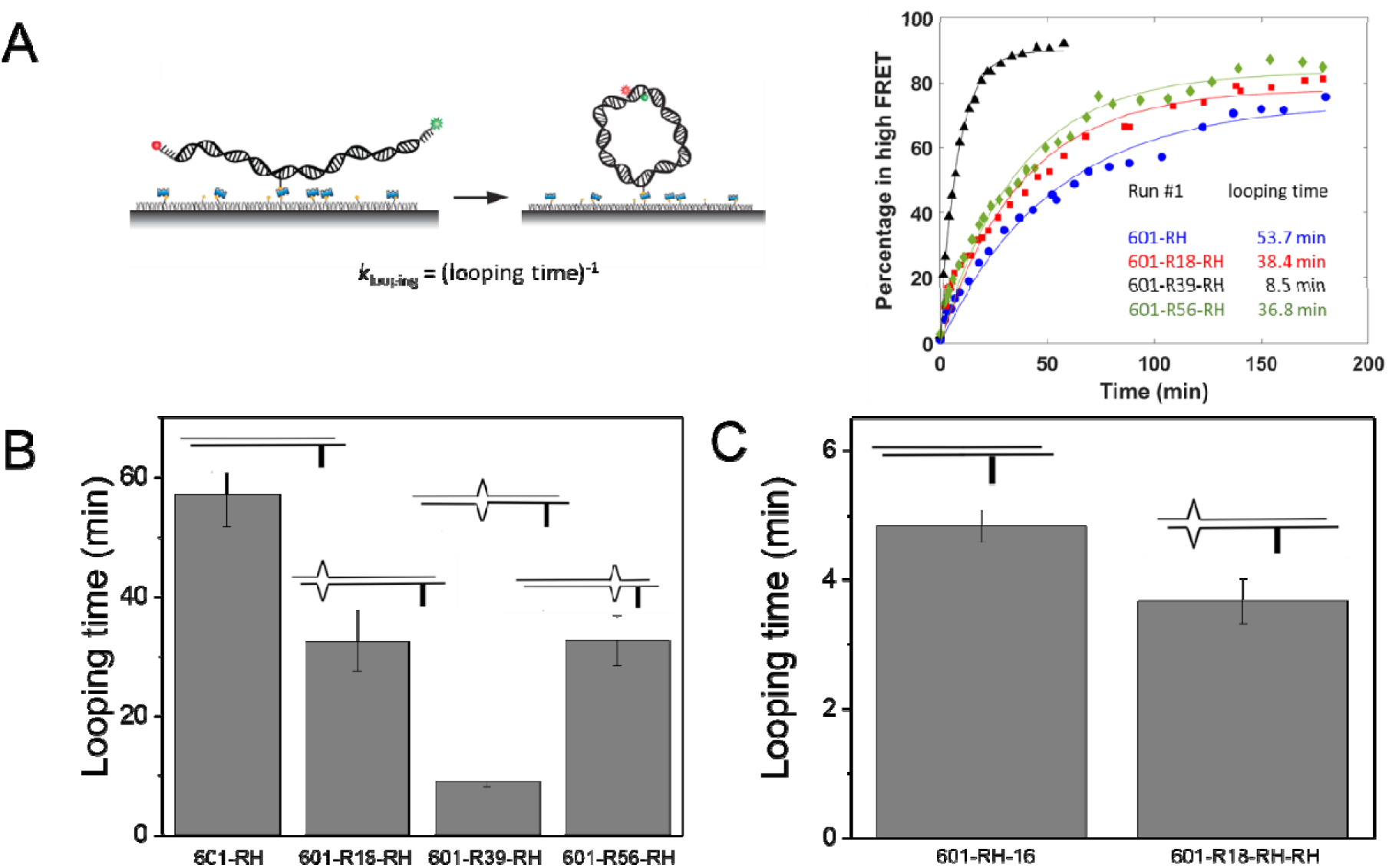
C-C mismatch enhances DNA flexibility. (**A**): Single molecule cyclization assay: The DNA construct with 10-nucleotide complementary sticky ends is immobilized on a PEG passivated imaging chamber. DNA looping is induced using the imaging buffer containing 1M NaCl followed by time course TIRF imaging. To calculate the looping time, the fraction of looped molecules (high FRET) as a function of time is fitted to an exponential function, *e*^-*t*/(*looping time*)^ (right panel for one run of experiments). (**B**, **C**) Fitted looping time for the right half of the 601 construct without and with mismatches (**B**) and with the biotin position being moved by 16 nt (**C**). Error bars represented the S.E.M with n = 3 technical replicates.

We measured the looping time of 4 DNA constructs corresponding to the right half of the 601 sequence (601-RH) with the addition of a C-C mismatch at the R18, R39 and R56 locations (601-R18-RH, 601-R39-RH and 601-R56-RH). As expected, we observed a dramatic decrease in looping time of the construct containing a mismatch (Fig. 5B). Adding a C-C mismatch reduced the looping time from 57 min to 32 min (601-R18-RH), 9 min (601-R39-RH), and 32 min (601-R56-RH). The reduction in apparent looping time was larger with the mismatch placing at the center (601-R39-RH) than toward the side of the RH fragment (601-R18-RH,601-R56-RH) likely because the looping measurement is more sensitive to the change in flexibility at the center of the DNA fragment.

The cyclizability of surface-tethered DNA constructs was shown to possess an oscillatory dependence on the position of the biotin tether ^41^. For example, moving the location of the biotin tether by half the helical repeat (∼ 5 bp) can lead to a large change in cyclization rate^41^, likely due to the preferred poloidal angle of a given DNA^42^ that determines whether the biotin is facing towards the inside of the circularized DNA, thereby hindering cyclization due to steric hindrance caused by surface tethering. We therefore performed control experiments to test the possibility that the observed higher cyclization rates of constructs with mismatches, as shown in Fig. 5B-C, is an artifact specific to the biotin tether location used. We created two additional constructs, 601-RH-16 and 601-R18-RH-16, which are identical to the 601-RH and 601-R18-RH constructs, respectively, except that the location of the biotin tether was moved to a thymine base that lies 16 nucleotides further towards the center of the molecule. We chose 16 nucleotides because it is about 1.5 times the helical repeat, and thus cyclization rates should be maximally different from those of the original 601-RH and 601-R18 constructs. Further, there was a thymine base present there to which the biotin could be conveniently attached. Side-by-side, we re-prepared the original 601-RH and 601-R18-RH constructs. We found that the overall looping rates of both the 601-RH-16 and 601-R18-RH-16 constructs were higher than those for the 601-RH and 601-R18-RH constructs, indicating that moving the biotin tether towards the center of the molecule increases looping rate (Fig. 5C). However, the 601-R18-RH-16 construct, which contains a mismatch, still looped faster than the 601-RH-16 construct without a mismatch (Fig. 5C). We thus conclude that the presence of the C-C mismatch makes the construct loop faster, and that this is not an artifact specific to the biotin tether location. As we will show next, the looping time for different mismatch types showed broadly similar behavior to that observed from DNA buckling experiments, further indicating that the mismatch effect is an intrinsic property.

### Effects of other mismatches on DNA bendability

There are eight different types of mismatches made from canonical DNA bases: A-A, T-T, C-C, G-G, G-A, C-A, C-T and G-T mismatches. We performed single molecule looping experiments of DNA containing a single mismatch introduced near the middle of the looping construct and determined the looping times for all eight constructs (Fig. 6). See Supplementary Materials for their sequences. We observed significant reduction in looping time compared to the intact DNA control (no mismatch) with the exception of G-containing mismatches (G-G, G-T and G-A mismatches). The C-containing mismatches (C-C, C-A and C-T) showed the largest reduction in looping times, suggesting that they make DNA most bendable. We also compared our looping times with the published measure of DNA bendability for the corresponding mismatched DNA where they quantified DNA buckling via FRET (*E*_FRET_ in Fig. 6)^6^. The two measures generally agree. However, we found sizable deviations from a linear relation for T-T and G-T mismatches. It is possible that the deviations may arise from sequence contexts, but we cannot exclude the possibility that the two assays measure slightly different aspects of DNA mechanics. Although performing fluorescence-force spectroscopy for non-C-C mismatches is beyond the scope of the current work, a testable prediction is that T-T and A-A mismatches as well as C-containing mismatches make a nucleosome mechanically more stable.

**Figure 6:**
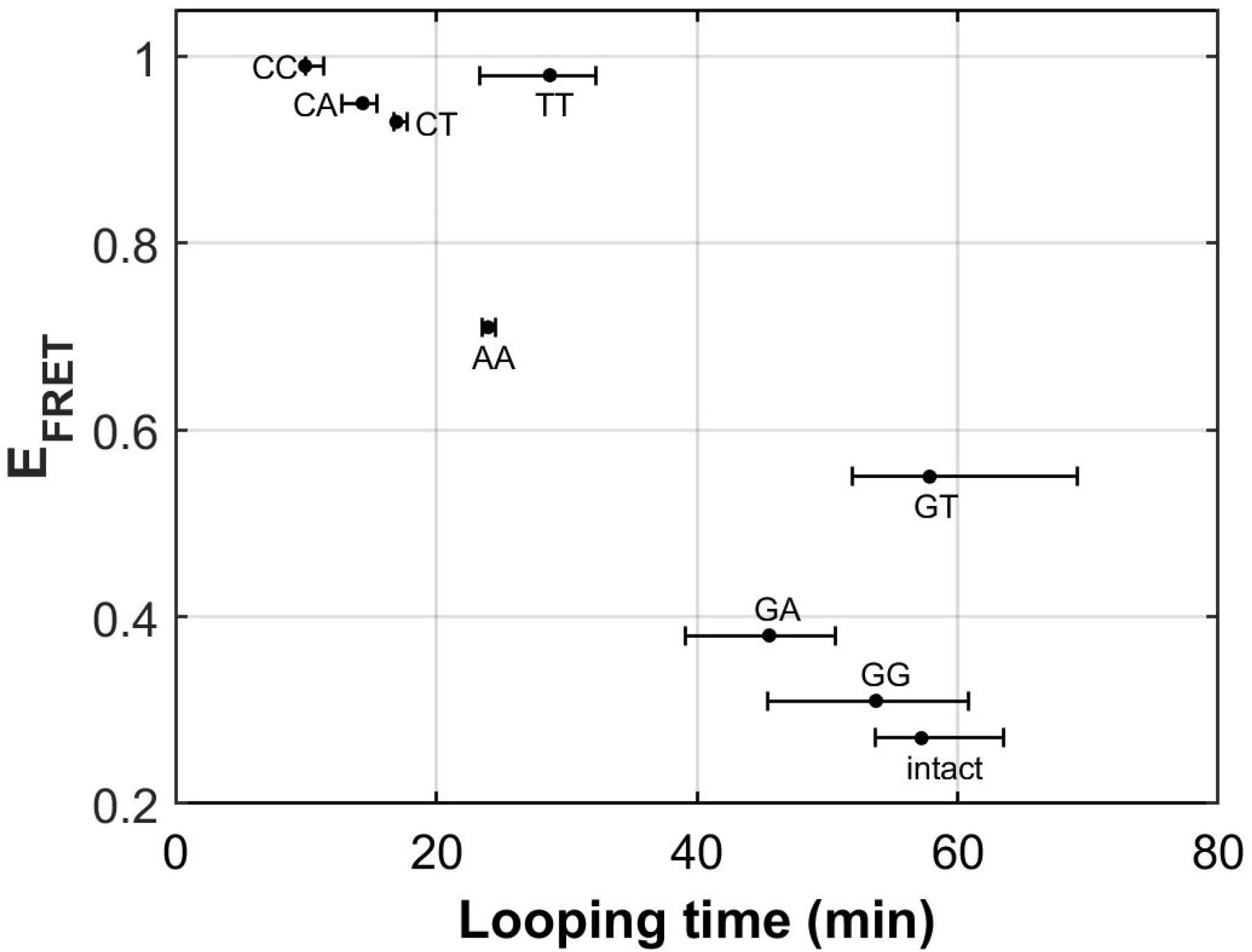
DNA flexibility enhancement is dependent on mismatch type. Looping times for DNA containing a single mismatch (one of eight types each) and an intact DNA without a mismatch. Also shown are ensemble FRET efficiencies (*E*_FRET_) from Ref. ^6^ as a measure of DNA buckling for the same type of mismatch.

### Yeast nucleosomes are less stable and more symmetrical than Xenopus nucleosomes in outer turn unwrapping

Next, we sought to examine how the source of histone proteins affects the mechanical stability of an intact nucleosome, i.e. without a mismatch. We reconstituted the 601 DNA construct with histone octamers of Xenopus and budding yeast. Note that all the data presented thus far on the effect of mismatches were obtained using Xenopus histones. Outer turn FRET probes on both sides ED1 and ED2 displayed slightly lower zero-force FRET values for yeast nucleosomes compared to Xenopus nucleosomes (Fig. 7), indicating that the DNA entry/exit may be more loosely bound on histone core for yeast nucleosomes. In contrast, the inner turn FRET probe showed similar zero-force FRET values for yeast and Xenopus nucleosomes. With pulling force applied, the stretching pattern for ED1 is similar for both nucleosomes while the strong side probe ED2 showed lower mechanical stability for yeast histones, with 40% of the molecules having unwrapping force of lower than 5 pN and the other 60% of the molecules being unwrapped by a force between 5-15 pN (Fig. 7A-B). The inner turn probe showed a stepwise unwrapping pattern with initial FRET reduction at less than 5 pN followed by stable FRET and a final unwrapping at a force higher than 20 pN (Fig. 7C). These observations for both outer turn and inner turn probes suggested that nucleosomes made with yeast histones are mechanically less stable, and unwrap less asymmetrically than nucleosomes made with Xenopus histones. Therefore, how DNA mechanics, determined by sequence, mismatch or chemical modification, is translated to nucleosome mechanics may be influenced by the histone core.

**Figure 7:**
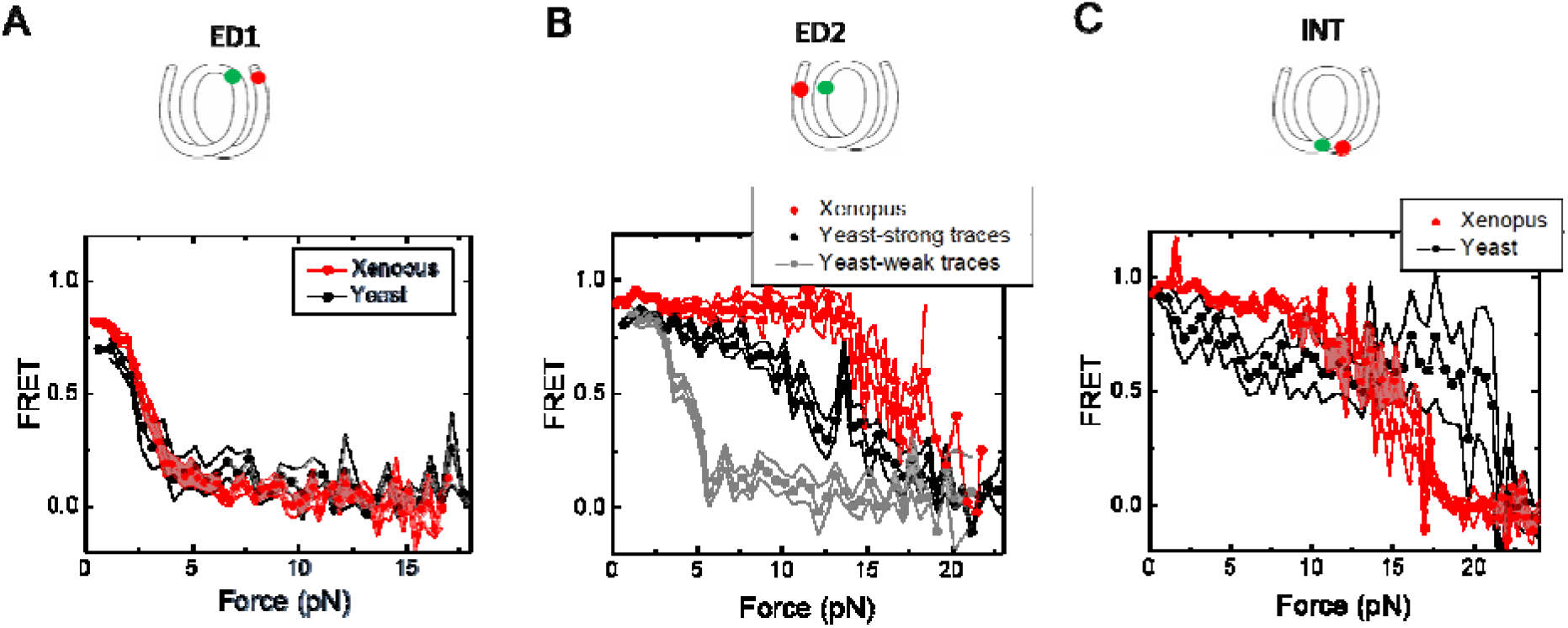
Unwrapping of yeast vs. xenopus reconstituted nucleosomes: Average of FRET vs. Force for nucleosomes reconstituted from xenopus (red) vs yeast (black and gray) histone proteins with DNA labeled by outer turn probes ED1 (**A**), ED2 (**B**) and inner turn probe INT (**C**). The error bars represent S.D. of n = 17 (Xenopus) and 5 (Yeast) nucleosomes with the ED1 probe (**A**), n = 20 (Xenopus), 6 (Yeast – strong) and 4 (Yeast-weak) nucleosomes with the ED2 probe (**B**), and n = 22 (Xenopus) and 6 (Yeast) nucleosomes with the INT probe (**C**), respectively.

## Discussion

Using the looping time of single molecule DNA cyclization as a measure of DNA bendability, we showed that a DNA mismatch can increase DNA bendability. Our results for the selected mismatches are consistent with previous studies on the effect of a mismatch on DNA conformational dynamics using other methods such as NMR ^7^ and the DNA Euler buckling assay ^6^. A possible explanation for the enhancement of DNA flexibility is the existence of a kink at the mismatch position on the DNA. If and how the mismatch type-dependent DNA mechanics affects the sequence-dependent mismatch repair efficiency in vivo, as recently determined in a high through study in *E. coli*^43^, remains to be investigated. Comparison of mismatch-type dependent DNA mechanics to population genetics data is challenging because mutation profiles reflect a combined outcome of mismatch-generation, mismatch repair and selection in addition to other mutational processes.

We observed the enhancement of mechanical stability of nucleosomes reconstituted from mismatch-containing DNA constructs. A defect making the system more stable may appear counterintuitive but given that the same mismatch can make DNA more flexible, our findings are in broad agreements with previous studies that showed positive correlation between DNA flexibility and nucleosome mechanical stability when DNA sequences or cytosine methylation was altered ^26,27^.

The 601 positioning sequence has TA-rich side that has four TA dinucleotides spaced with 10 bp periodicity and is more flexible than the TA-poor side ^26^. The 601 nucleosome is more stable on the TA-rich side ^26,35,44^, which we attributed to the ease with which the more bendable DNA stays sharply bent around the histone core even under unwrapping force. Here, we introduced a mismatch to the TA-poor side with the aim of achieving a large contrast in the background of rigid DNA. Indeed, the mismatch induced a ∼7-fold increase in the rate of DNA cyclization of the TA-poor side. This increase in DNA flexibility matches the ∼7 fold larger cyclization rate of the TA-rich side compared to the TA-poor side ^26^, suggesting that a single mismatch in the TA-poor side can symmetrize DNA flexibility of the 601 nucleosome. However, unlike flexibility symmetry achieved by TA repeats where which side unwraps at low forces became stochastic ^26^, when the flexibility symmetry was obtained via a mismatch in the TA-poor side, the TA-rich side remained mechanically stable, unwrapping at only high forces. This difference suggests that although the apparent flexibility is similar between DNA containing a mismatch vs a flexible sequence element, the mismatch does not have a global effect on the coordination of unwrapping of the two DNA ends.

The enhanced nucleosome mechanical stability we observed suggests that a mismatch will reduce nucleosomal DNA accessibility. The reduction in nucleosomal DNA accessibility would hinder the activity of the DNA mismatch repair machinery on nucleosomal DNA. An unrepaired mismatch leads to a point mutation which may be the source for genetic variation during evolution and cancer progression. In fact, previous observations showed that the frequency of single nucleotide polymorphism is higher near the nucleosome dyad for nucleosomes that are strongly positioned in vivo^21^. The higher frequency of substitutions in the nucleosomal DNA may be attributed to the difficulty of accessing the extra-stable nucleosomes. We also note that even without an enhanced stability, a mismatch within a nucleosome would be more difficult to detect for mismatch repair machineries compared to a mismatch in a non-nucleosomal DNA. Because mismatch repair machineries accompany the replisome, most of nascent mismatches may be detected for repair before nucleosome deposition. Therefore, the decrease in accessibility predicted based on our data here may be important only in rare cases a mismatch is not detected prior to the deposition of a nucleosome on the nascent DNA or in cases where a mismatch is generated via a non-replicative mechanism.

We chose the C-C mismatch for this work because a previous study showed that the C-C mismatch is one of the most flexible mismatches ^6^. If indeed more flexible elements in the DNA make a nucleosome mechanically stronger, as shown here for the C-C mismatch and previously for different sequences and cytosine modifications ^26,27^, we can predict that other mismatches, DNA lesions and alternative DNA structures such as DNA single strand damages, bulky adducts and R-loops ^45^ that alter DNA local flexibility would also change nucleosome mechanical stability accordingly. Future studies are needed to test this prediction.

We also tested if histones from different species and different histone variants can affect nucleosome stability under tension for an intact DNA without any mismatch. We observed a slightly lower zero-force FRET value for both sides with ED1 and ED2 for yeast nucleosomes compared to Xenopus nucleosomes. Under tension, we found that the outer turn of yeast nucleosomes could be unwrapped at a lower force than Xenopus nucleosomes, and that the unwrapping pattern is less asymmetric for ∼40% of nucleosomes formed on the 601 sequence. The crystal structure of the yeast nucleosome suggests that yeast nucleosome architecture is subtly destabilized in comparison with nucleosomes from higher eukaryotes^28^. Yeast histone protein sequences are not well conserved relative to vertebrate histones (H2A, 77%; H2B, 73%; H3, 90%; H4, 92% identities), and this divergence likely contributes to differences in nucleosome stability. Substitution of three residues in yeast H3 α3-helix (Q120, K121, K125) very near the nucleosome dyad with corresponding human H3.1/H3.3 residues (QK…K replaced with MP…Q) caused severe growth defects, elevated nuclease sensitivity, reduced nucleosome positioning and nucleosome relocation to preferred locations predicted by DNA sequence alone ^29^. The yeast histone octamer harboring wild type H3 may be less capable of wrapping DNA over the histone core, leading to reduced resistance to the unwrapping force for the more flexible half of the 601positioning sequence. Overall, our data suggest that how DNA mechanics, determined by sequence^26,41,46^, chemical modifications^27^ and mismatches^38,40^, is translated to nucleosome mechanics may be dependent on species-specific differences in histone sequence and post-translation modifications, which need to be examined in future studies.

Previous studies have showed that the artificial 601 sequence do not preferentially or strongly position the nucleosomes in vivo as expected ^47,48^. It is certainly desirable to perform mismatch-dependent and species-specific nucleosome mechanics studies using native sequences but the fluorescence-force spectroscopy data of the type we acquired in this study would be difficult to interpret unless the nucleosomes are formed at a well-defined position. We recently reported a native sequence from the yeast gene SWH1 that forms a nucleosome in vitro centered at the dyad position in vivo^33^.

While the enhancement of the mechanical stability of the nucleosome can potentially lead to the nucleosome’s decreased accessibility for the mismatch repair mechanism, which can account for the accumulation of single nucleotide polymorphisms near the nucleosome dyad, other functional implications of the enhanced mechanical stability cannot be precluded. For example, enhanced mechanical stability might shield DNA from transcription, which prevents the expression of genes that contain misincorporated nucleotides. Another opportunity for future studies is the fate of oligonucleosomes under tension when one of the nucleosomes contains a mismatch.

## Materials and Methods

### Preparation of labeled DNA constructs

Each strand of DNA in constructs for cyclization measurements was prepared by ligation of two shorter DNA fragments containing labeled Cy3, Cy5 and biotin as indicated in Supplementary Figure S1. Typically, the fragments were mixed at the ratio of 1:1.2:1.5 for the first, the helper, and the second fragments for ligation with T4 DNA ligase (NEB) following the manufacture manual. The ligation mixture was then loaded on a denaturing PAGE gel to run electrophoresis for purification. We cut and chop the top band which had the correct length and let the DNA diffuse to a buffer containing 10 mM Tris pH 8 and 50 mM NaCl. After purification, the two complementary strands were annealed by mixng at 1:1 molar ratio and heating to 90°C followed by slow cooling over 3-4 hours. The final DNA construct contained the 601 sequence and was flanked by a 14 bp spacer to biotin for surface tethering and 20 bp spacer connect to a 12 nts overhang for annealing to lambda DNA.

### Nucleosome preparation

Both *Xenopus laevis* and yeast histones were expressed in *E. coli*. Yeast histones were prepared in C. Wu’s lab at the National Institutes of Health as described ^49^. After purifying individual histone proteins, the histone octamers were prepared by denaturation-refolding and purification, according to standard procedures ^50^. Xenopus histone octamers were purchased from The Histone Source, Colorado State University. To prepare nucleosomes, 601 DNA templates were reconstituted with the recombinant histone octamer by step-wise salt-dialysis ^50^. The reconstituted nucleosome product was confirmed by an electrophoresis mobility shift assay for all experiments. Reconstituted nucleosomes were stored at 4°C in the dark, typically at concentrations of 100– 200 nM, and used within 4 weeks.

### Single molecule DNA cyclization Measurement

DNA fragments for cyclization measurement were immobilized on a PEG-coated microscope slide via biotin-neutravidin linkage. The fragments had complementary 10 nt 5’ overhang at either end, which permit looping via annealing. Cy3 and Cy5 were also present at the two 5’ ends, resulting in high FRET in the looped state. Measuring FRET allowed us to quantify the fraction of looped molecules as a function of time since introduction of a high salt buffered solution (10 mM Tris-HCl pH 8.0, 1 M NaCl, 0.5% w/v D-Glucose (Sigma), 165 U/ml glucose oxidase (Sigma), 2170 U/ml catalase (Roche) and 3 mM Trolox (Sigma). Approximately 2500 – 3500 molecules were quantified at each timestamp during the experiment, and three independent experiments were performed for each sequence (Supplemental Figure S2). The rate of loop formation was used as an operational measurement of DNA flexibility.

### Force-Fluorescence Spectroscopy Measurement

We followed the protocol for force-fluorescence spectroscopy measurement published previously ^26,27^. To construct the DNA tether for a Force-Fluorescence measurement, λ DNA was annealed to the reconstituted nucleosomes at one end, and to an oligonucleotide containing digoxigenin. The concentration of each element in the annealing reaction is 8 nM. During the experiment, the sample was diluted to 10 pM in nucleosome dilution buffer (10 mM Tris-HCl pH 8.0, 50 mM NaCl, 1 mM MgCl_2+_ or 1 mM spermine) for immobilization on the PEG coated microscope slide. To attach the micro beads for optical trapping to the DNA construct. we diluted 1 μm anti-digoxigenin-coated polystyrene beads (Polysciences) in nucleosome dilution buffer and added it to the imaging chamber for 30 minutes. The fluorescence-force data acquisition procedures include three following steps using a custom built setup according to ^30^. First, after trapping a bead, we determined the origin of the tether by stretching it in two opposite directions along x and y axis. Second, to spatially avoid beaching of the fluorophores, we displaced the trapped bead from its origin where the labeled nucleosome is located by 14 μm. To locate the exact position of the label nucleosomes for confocal acquisition of the fluorescence signal, we scan the confocal laser around the tether’s origin. Third, to apply the force on the tether, the nucleosome was stretched at a constant velocity of 455 nm/sec^26^

Fluorescence emission was recorded for 20 ms at each step during the stage movement by scanning the confocal excitation concurrently with the stage movement. Force-fluorescence data was obtained in imaging buffer (50 mM Tris-HCl pH 8, 50 mM NaCl, 1 mM MgCl_2_ or Spermine, 0.5 mg/ml BSA (NEB), 0.5 mg/ml tRNA (Ambion), 0.1% v/v Tween-20 (Sigma), 0.5% w/v D-Glucose (Sigma), 165 U/ml glucose oxidase (Sigma), 2170 U/ml catalase (Roche) and 3 mM Trolox (Sigma)). tRNA, which we normally include to reduce sticking of beads to the surface over the hours of single molecule experiments in a sealed chamber, was excluded in experiments with yeast-expressed nucleosomes because tRNA induced disassembly of nucleosomes assembled using yeast histones.

All single molecule measurements were performed at the room temperature.

## Acknowledgments

We thank Sergei Rudnizky for critical comments. This work was supported by the US National Institutes of Health (GM122569 to T.H. and GM125831 to C.W.) and by the National Science Foundation Physics Frontier Center program (PHY1430124). T.T.M.N is supported by the Cancer Early Detection Advanced Research Center (CEDAR) at Oregon Health and Science University and grants from the Department of Defense, Susan G. Komen Foundation, and Kuni Foundation. F.W. was supported by the NCI intramural research program. C.W. was a NIH Scientist Emeritus and a Senior Fellow of the Howard Hughes Medical Institute Janelia Research Campus. T.H. is an investigator with the Howard Hughes Medical Institute.

