## Supplementary Materials for "Dependence of Nucleosome Mechanical Stability on DNA Mismatches"

**Sequences for nucleosome reconstitution for the fleezers measurements – made by annealing the top and bottom strands which was constructed by ligation of two short fragments.** Light gray shades denote sequences in the outer turn of the nucleosome and dark gray shares denote sequences in the inner turn of the nucleosome. Green and red ‘T’s denote the labeling sites for donor (Cy3) and acceptor (Cy5) fluorophores, conjugation done via amino-dT for the ED1 construct. For labeling sites for the ED2 construct, we refer to Ngo et al^1^. Yellow highlights denote the positions of the mismatched bases. /idSp/ is the space added to prevent polymerization onto the 5’ overhang.

**601 original sequence (made by PCR)**

5’- /Biotin/TATA CGCGGCCGCC CTGGAGAATC CCGGTGCCGA GGCCGCTCAA TTGGTCGTAG ACAGCTCTAG CACCGCTTAA ACGCACG**T**AC GCGCTGTCCC CCGCGTTTTA ACCGCCAAGG GGATTACTCC CTAGTCTCCA GGCACGTGTC AGATATATAC ATCCTGT GCATGTATTG AACAGCGACC

ATAT GCGCCGGCGG GACCTCTTAG GGCCACGGCT CCGGCGAGTT AACCAGCATC TGTCGAGATC GTGGCGAATT TGCGTGCATG CGCGACAGGG GGCGCAAAAT TGGCGGTTCC CCTAATGAGG GATCAGAGGT CCGTGCACAG TCTATATATG **T**AGGACA CGTACATAAC TTGTCGCTGG/idSp/ TC CAGCGGCGGG – 5’

**R18 construct**

5’- /Biotin/TATA CGCGGCCGCC CTGGAGAATC CCGGTGCCGA GGCCGCTCAA TTGGTCGTAG ACAGCTCTAG CACCGCTTAA ACGCACG**T**AC GCGCTGTCCC CCGCGTTTTA ACCGCCAAGG GGATTACTCC CTAGTCTCCA GGCACGTGTC AGATATATAC ATCCTGT GCATGTATTG AACAGCGACC

ATAT GCGCCGGCGG GACCTCTTAG GGCCACGGCT CCGGCGAGTT AACCAGCATC TGTCGAGATC GTGGCGAATT TGCGTGCATG CGCGACAGGG GGCGCAAAAT TGGCGGTTCC CCTAATGAGG GATCAGAGGT CCGTGCACAC TCTATATATG **T**AGGACA CGTACATAAC TTGTCGCTGG/idSp/ TC CAGCGGCGGG – 5’

**R39 construct**

5’- /Biotin/TATA CGCGGCCGCC CTGGAGAATC CCGGTGCCGA GGCCGCTCAA TTGGTCGTAG ACAGCTCTAG CACCGCTTAA ACGCACG**T**AC GCGCTGTCCC CCGCGTTTTA ACCGCCAAGG GGATTACTCC CTAGTCTCCA GGCACGTGTC AGATATATAC ATCCTGT GCATGTATTG AACAGCGACC

ATAT GCGCCGGCGG GACCTCTTAG GGCCACGGCT CCGGCGAGTT AACCAGCATC TGTCGAGATC GTGGCGAATT TGCGTGCATG CGCGACAGGG GGCGCAAAAT TGGCGGTTCC CCTAATGACG GATCAGAGGT CCGTGCACAG TCTATATATG **T**AGGACA CGTACATAAC TTGTCGCTGG/idSp/ TC CAGCGGCGGG – 5’

**R56 construct**

5’- /Biotin/TATA CGCGGCCGCC CTGGAGAATC CCGGTGCCGA GGCCGCTCAA TTGGTCGTAG ACAGCTCTAG CACCGCTTAA ACGCACG**T**AC GCGCTGTCCC CCGCGTTTTA ACCGCCAAGG GGATTACTCC CTAGTCTCCA GGCACGTGTC AGATATATAC ATCCTGT GCATGTATTG AACAGCGACC

ATAT GCGCCGGCGG GACCTCTTAG GGCCACGGCT CCGGCGAGTT AACCAGCATC TGTCGAGATC GTGGCGAATT TGCGTGCATG CGCGACAGGG GGCGCAAAAT TCGCGGTTCC CCTAATGAGG GATCAGAGGT CCGTGCACAG TCTATATATG TAGGACA CGTACATAAC TTGTCGCTGG/idSp/ TC CAGCGGCGGG – 5’

**Sequences for looping measurements – made by annealing of the top and bottom strands which was synthesized by IDT**. Yellow highlights denote the locations of mismatched bases. Cyan highlights denote the location of biotin conjugated via dT.

```
601-RH
5’- /5Cy5/ACGGATTCTG TGTCCC CCGCGTT/iBiodT/TA ACCGCCAAGG GGATTACTCC CTAGTCTCCA GGCACGTGTC AGATATATAC ATCCTGT
                     ACAGGG GGCGCAAA       AT TGGCGGTTCC CCTAATGAGG GATCAGAGGT CCGTGCACAG TCTATATATG TAGGACA TGCCTAAGAC /5Cy3/ – 5’

601-R18-RH
5’- /5Cy5/ACGGATTCTG TGTCCC CCGCGTT/iBiodT/TA ACCGCCAAGG GGATTACTCC CTAGTCTCCA GGCACGTGTC AGATATATAC ATCCTGT
                     ACAGGG GGCGCAAA       AT TGGCGGTTCC CCTAATGAGG GATCAGAGGT CCGTGCACAC TCTATATATG TAGGACA TGCCTAAGAC /5Cy3/ – 5’

601-R39-RH
5’- /5Cy5/ACGGATTCTG TGTCCC CCGCGTT/iBiodT/TA ACCGCCAAGG GGATTACTCC CTAGTCTCCA GGCACGTGTC AGATATATAC ATCCTGT
                     ACAGGG GGCGCAAA       AT TGGCGGTTCC CCTAATGACG GATCAGAGGT CCGTGCACAG TCTATATATG TAGGACA TGCCTAAGAC /5Cy3/ – 5’

601-R56-RH
5’- /5Cy5/ACGGATTCTG TGTCCC CCGCGTT/iBiodT/TA ACCGCCAAGG GGATTACTCC CTAGTCTCCA GGCACGTGTC AGATATATAC ATCCTGT
                     ACAGGG GGCGCAAA       AT TCGCGGTTCC CCTAATGAGG GATCAGAGGT CCGTGCACAG TCTATATATG TAGGACA TGCCTAAGAC /5Cy3/ - 5’

601-RH-16
5’- /5Cy5/ACGGATTCTG TGTCCC CCGCGTTTTA ACCGCCAAGG GGA/iBiodT/TACTCC CTAGTCTCCA GGCACGTGTC AGATATATAC ATCCTGT
                     ACAGGG GGCGCAAAAT TGGCGGTTCC CCTA       ATGAGG GATCAGAGGT CCGTGCACAG TCTATATATG TAGGACA TGCCTAAGAC /5Cy3/ – 5’

601-R18-RH-16
5’- /5Cy5/ACGGATTCTG TGTCCC CCGCGTTTTA ACCGCCAAGG GGA/iBiodT/TACTCC CTAGTCTCCA GGCACGTGTC AGATATATAC ATCCTGT
                     ACAGGG GGCGCAAAAT TGGCGGTTCC CCTA       ATGAGG GATCAGAGGT CCGTGCACAC TCTATATATG TAGGACA TGCCTAAGAC /5Cy3/ – 5’

R40-RH-TT
5’- /5Cy5/ACGGATTCTG TGTCCC CCGCGTT/iBiodT/TA ACCGCCAAGG GGATTACTCC CTAGTCTCCA GGCACGTGTC AGATATATAC ATCCTGT
                     ACAGGG GGCGCAAA       AT TGGCGGTTCC CCTAATGTGG GATCAGAGGT CCGTGCACAG TCTATATATG TAGGACA TGCCTAAGAC /5Cy3/ – 5’


R40-RH-AA
5’- /5Cy5/ACGGATTCTG TGTCCC CCGCGTT/iBiodT/TA ACCGCCAAGG GGATTACACC CTAGTCTCCA GGCACGTGTC AGATATATAC ATCCTGT
                     ACAGGG GGCGCAAA       AT TGGCGGTTCC CCTAATGAGG GATCAGAGGT CCGTGCACAG TCTATATATG TAGGACA TGCCTAAGAC /5Cy3/ – 5’


R39-RH-CT
5’- /5Cy5/ACGGATTCTG TGTCCC CCGCGTT/iBiodT/TA ACCGCCAAGG GGATTACTCC CTAGTCTCCA GGCACGTGTC AGATATATAC ATCCTGT
                     ACAGGG GGCGCAAA       AT TGGCGGTTCC CCTAATGATG GATCAGAGGT CCGTGCACAG TCTATATATG TAGGACA TGCCTAAGAC /5Cy3/ – 5’


R39-RH-CA
5’- /5Cy5/ACGGATTCTG TGTCCC CCGCGTT/iBiodT/TA ACCGCCAAGG GGATTACTCC CTAGTCTCCA GGCACGTGTC AGATATATAC ATCCTGT
                     ACAGGG GGCGCAAA       AT TGGCGGTTCC CCTAATGAAG GATCAGAGGT CCGTGCACAG TCTATATATG TAGGACA TGCCTAAGAC /5Cy3/ – 5’


R41-RH-GG
5’- /5Cy5/ACGGATTCTG TGTCCC CCGCGTT/iBiodT/TA ACCGCCAAGG GGATTAGTCC CTAGTCTCCA GGCACGTGTC AGATATATAC ATCCTGT
                     ACAGGG GGCGCAAA       AT TGGCGGTTCC CCTAATGAGG GATCAGAGGT CCGTGCACAG TCTATATATG TAGGACA TGCCTAAGAC /5Cy3/ – 5’


R41-RH-GA
5’- /5Cy5/ACGGATTCTG TGTCCC CCGCGTT/iBiodT/TA ACCGCCAAGG GGATTAATCC CTAGTCTCCA GGCACGTGTC AGATATATAC ATCCTGT
                     ACAGGG GGCGCAAA       AT TGGCGGTTCC CCTAATGAGG GATCAGAGGT CCGTGCACAG TCTATATATG TAGGACA TGCCTAAGAC /5Cy3/ – 5’


R41-RH-GT
5’- /5Cy5/ACGGATTCTG TGTCCC CCGCGTT/iBiodT/TA ACCGCCAAGG GGATTATTCC CTAGTCTCCA GGCACGTGTC AGATATATAC ATCCTGT
                     ACAGGG GGCGCAAA       AT TGGCGGTTCC CCTAATGAGG GATCAGAGGT CCGTGCACAG TCTATATATG TAGGACA TGCCTAAGAC /5Cy3/ – 5’
```

**Supplementary Figures**

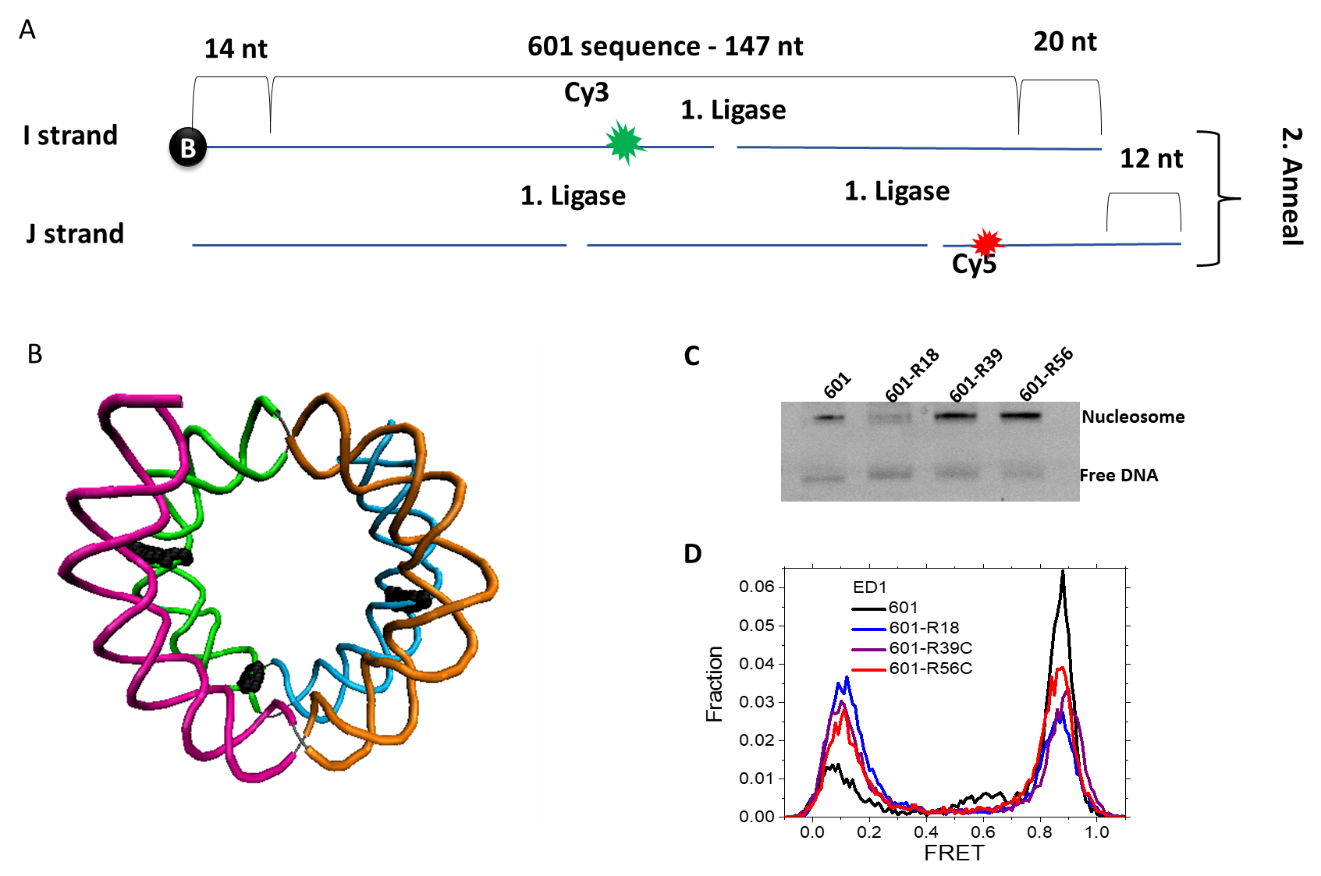

**Supplementary Figure S1**: Nucleosome preparation

(A) Scheme of the DNA template prepared by ligation of short, labeled oligos.
(B) DNA structure marking three sites of mismatch insertion (R56, R39 and R18, running from left to right)

**(C)** Migration of the 601 nucleosome mismatch containing nucleosomes on 5% native PAGE.

**(D)** FRET histogram of the 601 nucleosome mismatch containing nucleosomes with ED1 labeling scheme. The low FRET peak contains nucleosomes without a fluorescently active acceptor/

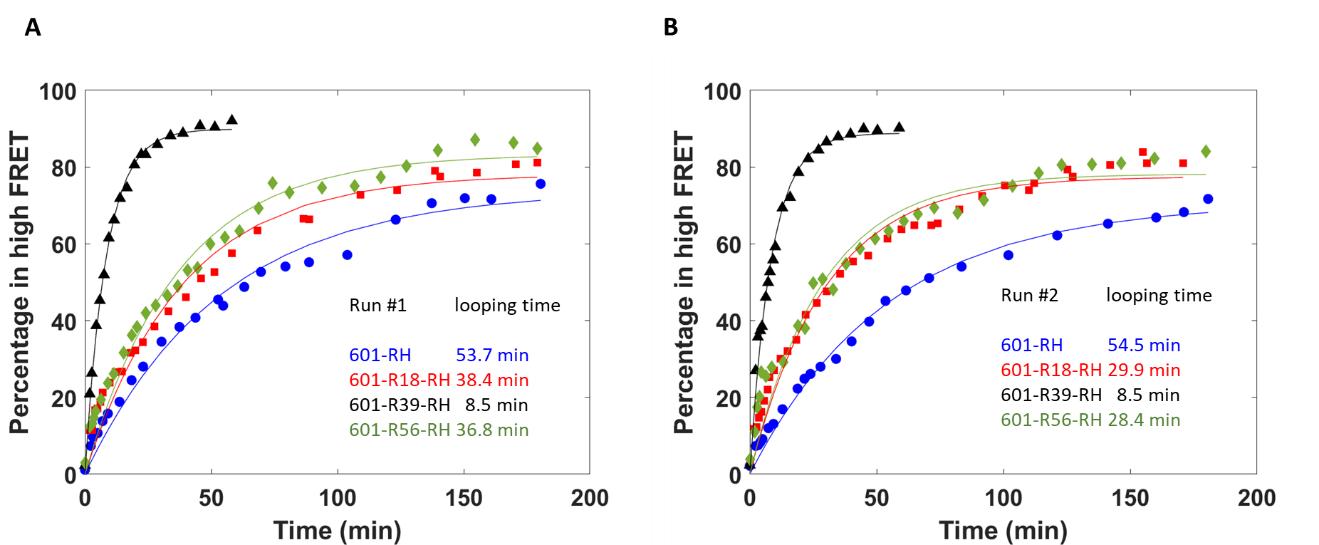

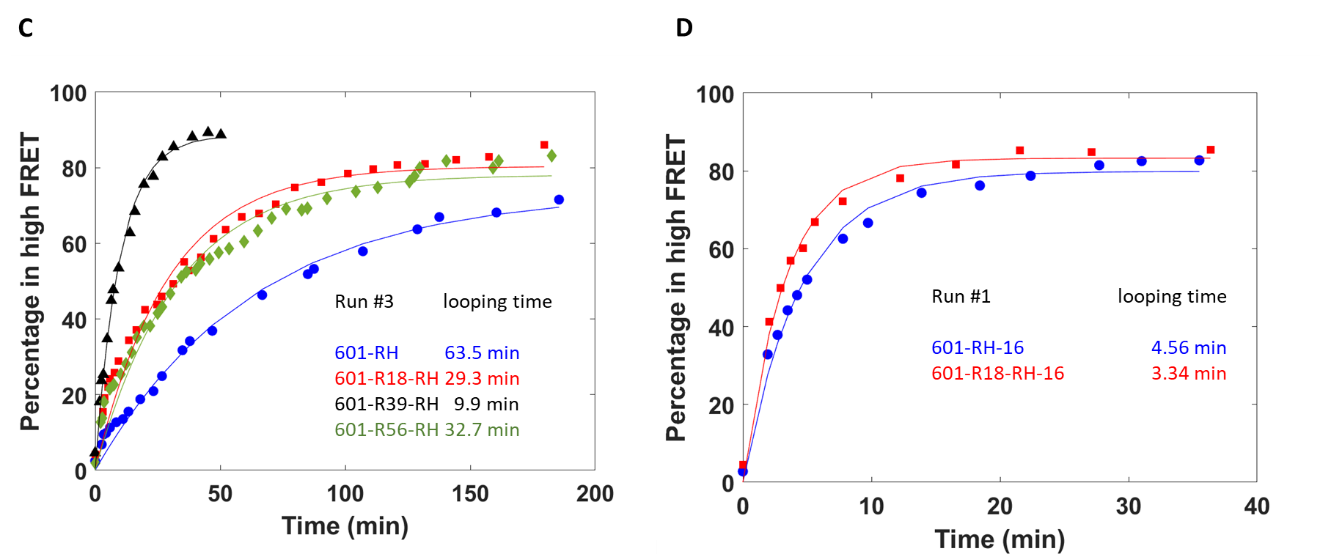

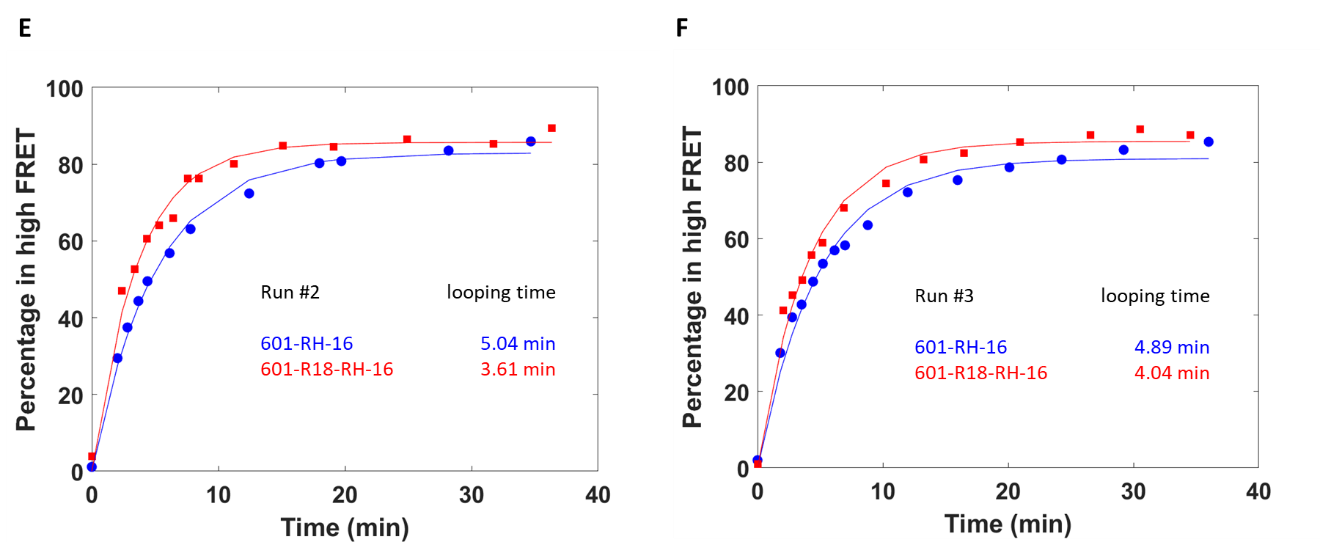

**Supplementary Figure S2**: Single molecule cyclization time course quantification. Each run has four times courses of looped fraction (percentage in high FRET population) vs time. Approximately 2500 – 3500 molecules were quantified at each timestamp during the experiment, and three independent experiments (run 1, run 2 and run 3) were performed for each sequence.

(A) - (C) Fraction of DNA molecules in high FRET over time for the 601 sequences with a C-C mismatch. Run 1 in panel A is also shown in Figure 5.

(D) - (F) Fraction of DNA molecules in high FRET over time for the 601 sequences with a C-C mismatch and biotin moved 16 nucleotides toward the center of the construct.
